# Superbubbles, Ultrabubbles and Cacti

**DOI:** 10.1101/101493

**Authors:** Benedict Paten, Adam M Novak, Erik Garrison, Glenn Hickey

## Abstract

A superbubble is a type of directed acyclic subgraph with single distinct source and sink vertices. In genome assembly and genetics, the possible paths through a superbubble can be considered to represent the set of possible sequences at a location in a genome. Bidirected and biedged graphs are a generalization of digraphs that are increasingly being used to more fully represent genome assembly and variation problems. Here we define snarls and ultrabubbles, generalizations of superbubbles for bidirected and biedged graphs, and give an efficient algorithm for the detection of these more general structures. Key to this algorithm is the cactus graph, which we show encodes the nested decomposition of a graph into snarls and ultrabubbles within its structure. We propose and demonstrate empirically that this decomposition on bidirected and biedged graphs solves a fundamental problem by defining genetic sites for any collection of genomic variations, including complex structural variations, without need for any single reference genome coordinate system. Furthermore, the nesting of the decomposition gives a natural way to describe and model variations contained within large variations, a case not currently dealt with by existing formats, e.g. VCF.

## 1 Introduction

Graphs are used extensively in biological sequence analysis, where they are often used to represent uncertainty about, or ensembles of, potential nucleotide sequences. Several subtypes have become especially prominent for sequence representation, in particular the De Bruijn graph [4,13], the string graph [10], the breakpoint graph [1,14] and the bidirected graph (aka variation graph or sequence graph) [6,9].

In the context of de novo sequence assembly several characteristic types of subgraph are recognised, in particular the *bubble* [16], a pair of paths that start and end at common source and sink nodes but are otherwise disjoint. In the context of sequence analysis, a bubble can represent a potential sequencing error or a genetic variation within a set of homologous molecules. An efficient algorithm for bubble detection was proposed by [2].

A generalization of the notion of a bubble, the superbubble is a more complex subgraph type in which a set of (not necessarily disjoint) paths start and end at common source and sink nodes. This problem was initially proposed by [11], who gave a quadratic solution. [3] recently provided a linear time algorithm for superbubbles on directed acyclic graphs (DAGs). This result, when paired with a previous linear time transformation of the problem of superbubbles on directed graphs to superbubbles on DAGS [15], yields a linear cost solution for computing superbubbles on digraphs. For a review of superbubbles and their use in sequence analysis see [8]. In this paper we generalize the idea of superbubble to the more general case of a bidirected graph, connect a slight generalization of the superbubble, which we call the ultrabubble, and show how it relates to the decomposition of the graph into 2- and 3-edge connected components.

## 2 Methods

### 2.1 Directed, Bidirected and Biedged Graphs

A *bidirected graph D* = (*V_D_*,*E_D_*) is a graph in which each endpoint of every edge has an independent orientation (denoted either “left” or “right”), indicating if the endpoint is incident with the left or right *side* of the given vertex. The sides of *D* are therefore the set *V_D_* × {*left*, *right*}, and each edge in *E_D_* is a pair set of two sides (Fig.1). We say for all *x* ∈ *V_D_*, (*x*,*left*) and (*x*, *right*) are *opposite sides*.

Any digraph is a special case of a bidirected graph in which each edge connects a left and a right side (by convention we here consider the right side to be the outgoing side and the left side the incoming side, so that the conversion from a digraph to a bidirected graph is determined; see Fig. 1).

**Fig. 1.**
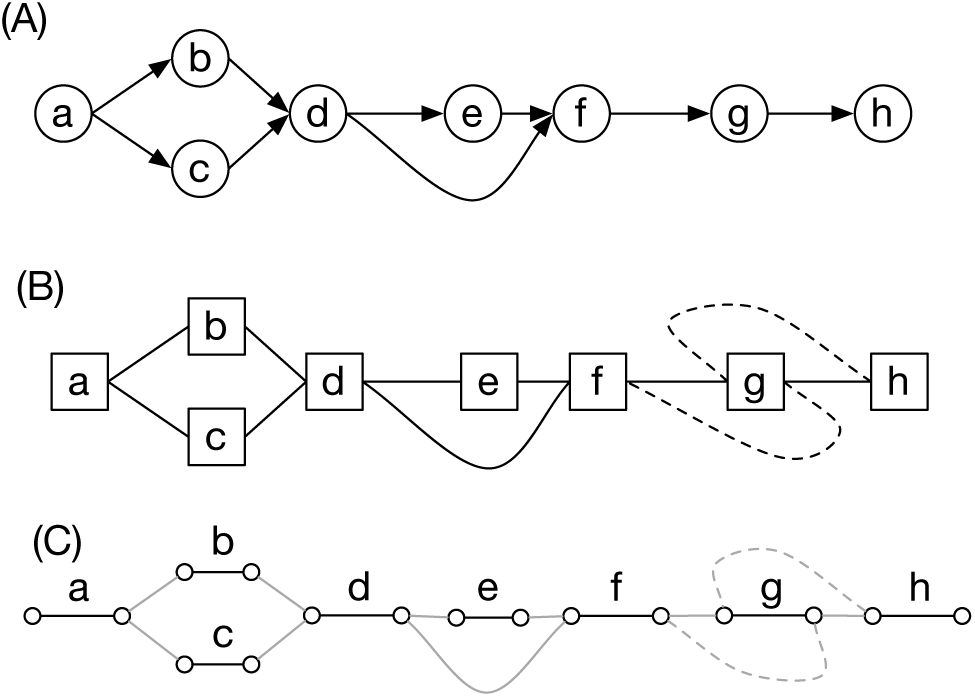
(A) A digraph. (B) A bidirected graph. Each node is drawn as a box and the orientation for each edge endpoint is indicated by the connection to either the left or right side of the node. The graph excluding the dotted edges is the equivalent bidirected graph for the digraph in (A); the dotted edges encode an inversion that cannot be expressed in the digraph representation. (C) A biedged graph equivalent to the bidirected graph shown in (B).

A *biedged graph* is a graph with two types of edges: *black edges* and *grey edges*, such that each vertex is incident with at most one black edge (Fig. 1(C)).

For any bidirected graph *D* there exists an equivalent biedged graph *B*(*D*) = (*V_B_*_(*D*)_, *E_B_*_(*D*)_) where:

- *V_B_*_(*D*)_ = *V_D_* × {*left*, *right*}, the sides of *V_D_*.
- *E_B_*_(*D*)_ = *S_B_*_(*D*)_ ∪ *E_D_*, where *E_D_* are the grey edges,
- and *S_B_*_(*D*)_ = {{(*x*, *left*), (*x*, *right*)}|*x* ∈ *V_D_*} are the black edges.

For a vertex *x* ∊ *V_B_*_(*D*)_ we use the notation *x*̂ to denote the opposite side to *x*, e.g. for *x* = (*x*′, *left*) ∈ *V_B_*_(*D*)_, *x*̂ = (*x*′,*right*).

Clearly the bidirected and biedged representations are essentially equivalent, and the choice to use either one is largely a stylistic consideration. For the remainder of this paper we will mostly use the biedged representation. As any digraph is a special case of a bidirected graph and any bidirected graph has a equivalent biedged graph, so any digraph has an equivalent biedged graph.

### 2.2 Directed Walks on Biedged and Bidirected Graphs

A directed walk on a bidirected graph is a walk that at each visited vertex exits the opposite side to that which it enters. On a biedged graph a directed walk is equivalent to a walk that alternates between black and grey edges. A bidirected or biedged graph is acyclic if it contains no directed cycles.

These definitions are a generalization of a directed walk on a digraph. In a bidirected representation of a digraph all edges in a directed walk are all left-to-right or all right-to-left. A directed walk on a general bidirected (or biedged) graph can mix these two types and additionally include edges that do not alternate the orientation of their endpoints (e.g. left-right and right-right and left-left edges).

Given these generalizing relationships, clearly a digraph **D** is acyclic iff *B*(**D**) is acyclic. Note that any acyclic biedged graph can also be converted into an equivalent directed acyclic graph (DAG):

#### Lemma 1.

*For any acyclic biedged graph B*(*D*) *there exists an isomorphic biedged graph B*(***D***) *such that **D** is a DAG.*

*Proof*. Use a depth first search (DFS) beginning at side *x* to label the sides of *B*(*D*) either ‘red’ or ‘white’: If *x* is not already labelled then label *x* red and *x*̂ white. For each grey edge incident with *x*̂, if the connected side is not labeled, label the connected side red and continue recursively via DFS. In this way all the sides in the connected component containing *x* will be labeled in a single DFS. If during the recursion the connected side encountered is already labelled then it must be labeled red, else there would exist a cycle in the DFS, a contradiction. Use the labelling to create *B*(**D**), isomorphic to *B*(**D**) but replacing the orientation of the sides so that each side labeled white is a left side and each side labeled red is a right side. All edges in *B*(**D**) connect a left and a right side.

### 2.3 Superbubbles, Snarls and Ultrabubbles

Repeating the definition from [11], any pair of distinct vertices (*x*,*y*) in a digraph **D** is called a *superbubble* (Fig. 2(A)) if:

- *reachability*: *y* is reachable from *x*.
- *matching*: The set of vertices, *X*, reachable from *x* without passing through *y* is equal to the set of vertices from which *y* is reachable without passing through *x* (passing through here means to enter and then exit a vertex on the path).
- *acyclicity*: The subgraph induced by *X* is acyclic.
- *minimality*: No vertex in *X* other than *y* forms a pair with *x* that satisfies the criteria defined above, and similarly for *y*.

We call the subgraph induced by *X* the *superbubble subgraph*.

To generalize superbubbles for biedged graphs we introduce the notion of a snarl, a minimal subgraph in a biedged graph whose vertices are at most 2-black-edge-connected (2-BEC) to the remainder of the graph (two vertices in a biedged graph are *k*-black-edge-connected (*k*-BEC) if it takes the deletion of at least *k* black edges to disconnect them). In a biedged graph *B*(*D*) a pair set of distinct, non-opposite vertices {*x*, *y*} are a *snarl* (Fig. 2(B)) if:

- *seperable*: The removal of the black edges incident with *x* and *y* disconnects the graph, creating a *separated component X* containing *x* and *y* and not *x*̂ and *y*̂.
- *minimality*: No node *z* in *X* exists such that {*x*, *z*} fulfils the above criteria, and similarly for *y*.

**Fig. 2.**
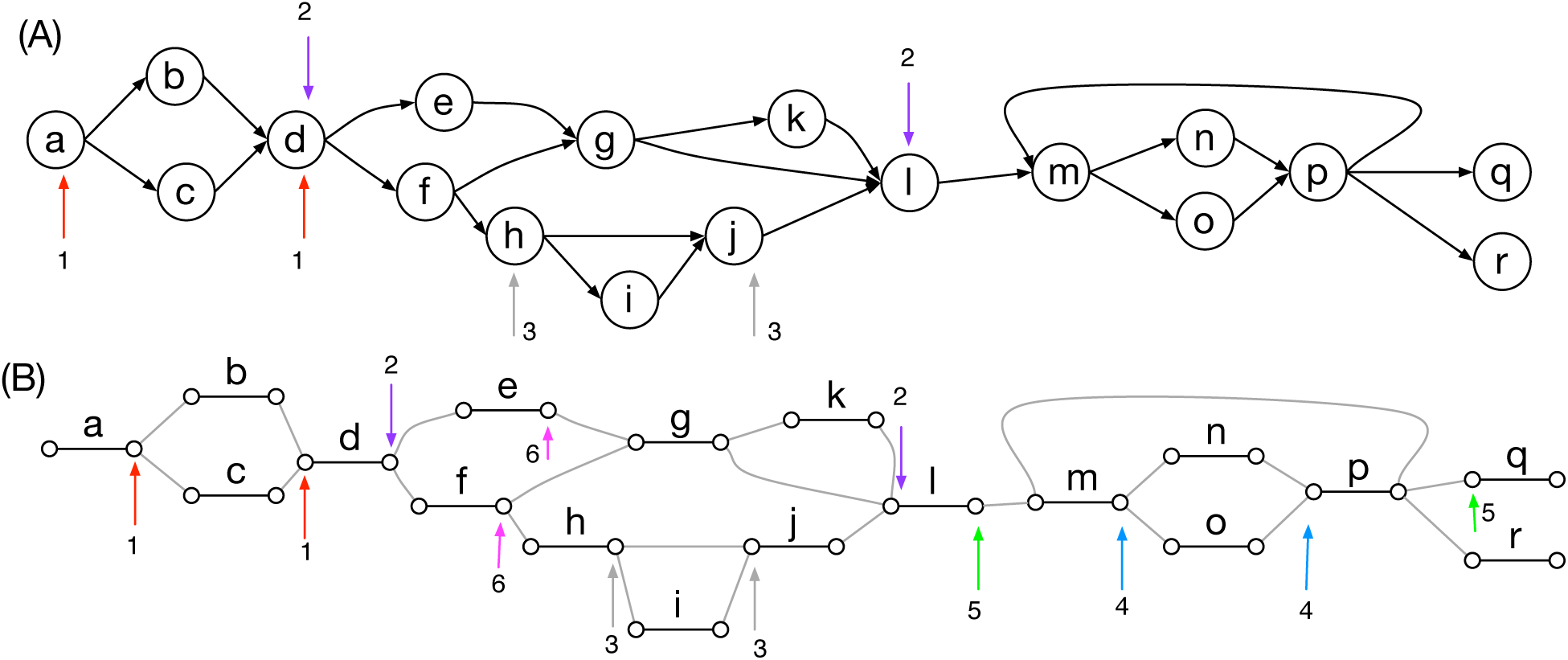
(A) Superbubbles in a digraph. The superbubbles are indicated by pairs of numbered arrows. (B) A biedged graph representation of the digraph in (A). The ultrabubbles are illustrated, as are two snarls that are not ultrabubbles (of several; pairs 5 and 6, whose separated components contain cycles).

We call a vertex not incident with a grey edge a *tip* [16]. In a biedged graph *B*(*D*) a snarl is an *ultrabubble* if its separated component is acyclic and contains no tips.

The following shows that a superbubble in a digraph is an ultrabubble in the equivalent biedged graph.

#### Lemma 2.

*For any superbubble* (*x*, *y*) *in a digraph **D***, *the pair set* {*x*′ = (*x*, *right*), *y*′ = (*y*, *left*)} *is an ultrabubble in B*(***D***).

*Proof.* Let *d* and *e* be the black edges incident with *x*′ and *y*′, respectively, and let *X* be the superbubble subgraph of (*x*, *y*).

We start by proving that {*x*′, *y*′} satisfies the separable criteria. As *y* is reachable from *x* by definition there exists a directed path in *B*(**D**) between *x*′ (the right side of *x*) and *y*′ (the left side of *y*) that excludes *d* and *e*. After the deletion of these black edges *x*′ and *y*′ therefore remain connected. If the separable criteria is not satisfied the deletion of *d* and *e* must therefore not disconnect *x*′ and *y*′ from either or both 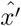 and 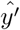, without loss of generality assume *x*′ (and therefore *y*′) remains connected to 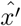.

If 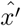 is on a directed walk from *x*′ that excludes *d* then the addition of *d* to this walk defines a directed cycle in *B*(**D**). As all nodes reachable from *x* are in the separated component *X*, the existence of this cycle in *B*(**D**)) implies the existence of a corresponding directed cycle in *X*, a contradiction.

If there exists a non-directed walk from *x*′ to 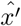 then let *z*′ be the last node on the walk from *x*′ such that the subwalk between *x*′ and *z*′ is a directed walk. By definition, there exists directed walk from *z*′ to *y*′. The next node on the walk from *x*′ to 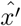 after *z*′ is, by definition, not reachable from *x*′ but *y*′ must be reachable from this node. This implies a contradiction of the matching criteria for the corresponding nodes in *X*.

We have therefore established that {*x*′, *y*′} fufills the seperable criteria. We have already established that iff a digraph is acyclic its equivalent biedged graph is acyclic, therefore the seperated component of {*x*′, *y*′} is acyclic. As every node in *X* is both reachable from *x* and on a path from *y*, the separated component clearly contains no tips.

It remains to prove that {*x*′, *y*′} fufills the minimality criteria. If {*x*′, *y*′} do not satisfy the minimality criteria without loss of generality there exists a node *z*′ in the separated component of {*x*′, *y*′} such that {*x*′, *y*′} are separable. It follows that all directed paths from *x*′ to *y*′ that exclude *d* and *e* visit *z*′, and for the node *z* in **D** contained in *z*′, (*x*, *z*) fulfills (clearly) all the superbubble criteria, a contradiction.

### 2.4 Cactus Graphs

A cactus graph is a graph in which any two vertices are at most two-edge connected [7]. In a cactus graph each edge is part of at most one simple cycle, and therefore any two simple cycles intersect at most one vertex.

For a graph *G* = (*V_G_*, *E_G_*) let *G′* = (*V_G′_*, *E_G′_*) be a multigraph created by *merging* subsets of the vertices, such that:

- *V_G′_* is a partition of *V_G_*,
- *E_G′_* = {{*a_G′_*(*x*),*a_G′_*(*y*)}|{*x*,*y*} ∈ *E_G_*} is a multiset.

where *a_G′_*: *V_G_* → *V_G′_* is a graph homomorphism that maps each vertex in *V_G_* to the set in *V_G′_* that contains it.

Merging all equivalence classes of 3-edge connected (3-EC) vertices in a graph results in a cactus graph [12].

For a biedged graph *B*(*D*) let *C*(*D*) be the cactus graph created by first contracting all the grey edges in *B*(*D*) then for each equivalence class of 3-EC vertices in the resulting graph merging together the vertices within the equivalence class (Fig. 3(A-C)). As with *G*′ and *G*, *V_C_*_(*D*)_ is a partition of the vertices of *V_B_*_(*D*)_, and *E_C_*_(*D*)_ = *{{*a_C_*_(*D*)_(*x*),*a_C_*_(*D*)_(*y*)}|{*x*,*y*} ∈ *S_B_*_(*D*)_}* is a multiset.

**Fig. 3.**
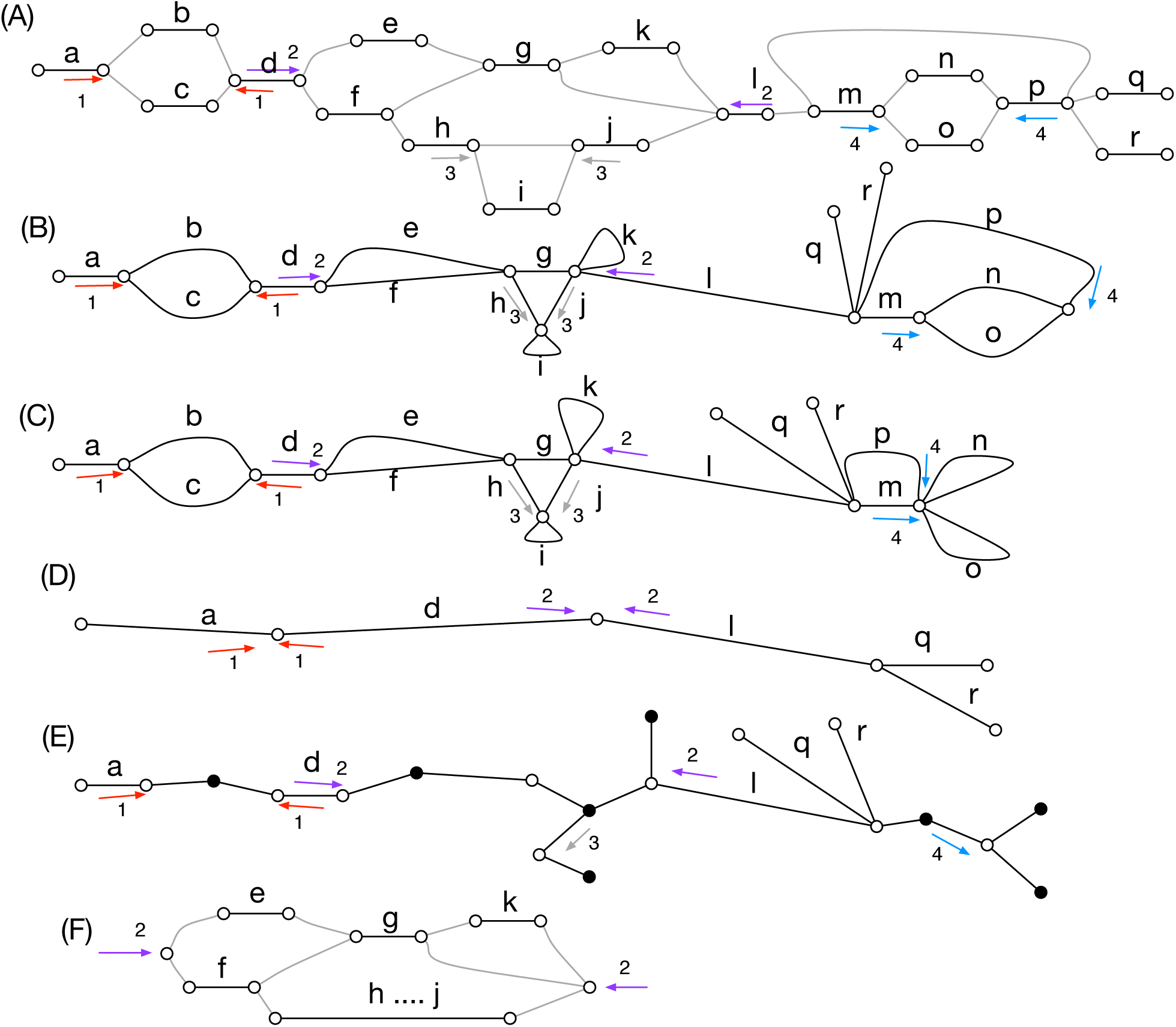
(A) A biedged graph *B*(*D*) with ultrabubbles indicated by pairs of numbered arrows. (B) The graph in (A) after contracting the grey edges. (C) The cactus graph *C*(*D*) for *B*(*D*). (D) The bridge forest *D*(*D*). (E) The cactus tree *T*(*D*). (F) A net for the number 2 bridge pair in (A). The projection of chain pairs in *B*(*D*) to the other graphs is shown using the numbered arrows, with the arrows drawn along the projecting black edge incident with the projected vertex.

For a vertex *x* ∈ *V_B_*_(*D*)_ we call *a_C_*_(*D*)_ its *projection* (in *C*(*D*)). Similarly for a set of vertices *X* ⊂ *V_B_*_(*D*)_ we call {*a_C_*_(*D*)_(*x*)|*x* ∈ *X*} the projection of *X* (in *C*(*D*)). Let *b_C_*_(*D*)_(*x*) = {*a_C_*_(*D*)_(*x*), *a_C_*_(*D*)_(*x* ̂)}, which is the projection of the black edge incident with *x* in *C*(*D*).

Appendix 1 gives lemmas that make explicit the relationship between the edge connectivity of vertices in *B*(*D*) and *C*(*D*), and which we use to prove the relationship between the snarls of *B*(*D*) and *C*(*D*).

### 2.5 Snarls and Cacti

A pair set of distinct vertices {*x*, *y*} in *B*(*D*) are a *chain pair* if they project to the same vertex in *C*(*D*) and their incident black edges project to the same simple cycle in *C*(*D*) (e.g. grey arrows and cyan arrows in Fig. 3(C)). A cyclic sequence of chain pairs within the same simple cycle in *C*(*D*) and ordered according to the ordering of this simple cycle is a *(cyclic) chain*. Contiguous chain pairs in a chain share two opposite sides of a black edge in *B*(*D*).

For a cactus graph *C*(*D*), the graph *D*(*D*) resulting from contracting all the edges in simple cycles in *C*(*D*) is a called a *bridge forest* (Fig. 3(D)).

A pair set of distinct vertices {*x*, *y*} in *B*(*D*) are a *bridge pair* if they project to the same vertex in *D*(*D*) and both their incident black edges are bridges (e.g. pairs of arrows numbered 1 and 2 in Fig. 3(D)). A maximum sequence of bridge pairs within *D*(*D*) connected by incident nodes with degree two is an *(acyclic) chain*. As with chain pairs, contiguous bridge pairs in a chain share two opposite sides of a black (bridge) edge in *B*(*D*).

#### Theorem 1.

*The set of snarls in B*(*D*) *is equal to the union of chain pairs and bridge pairs*.

*Proof*. Follows from Lemmas 12 and 13 given in Appendix 2.

Given Theorem 1 to calculate the set of snarls for a given biedged graph it is sufficient to calculate the cactus graph to give the set of snarls that map to chain pairs and the bridge forest to calculate the set of snarls that map to bridge pairs. Constructing a cactus graph of the type described for a biedged graph is linear in the size of the biedged graph (using the algorithm described in [12]), and clearly the cost of then calculating the bridge forest from the cactus graph is similarly linear. The number of chain pairs is clearly linear in the size of the biedged graph, however, the number of bridge pairs is potentially quadratic in the number of bridge pairs, so enumerating these latter snarls has potentially worst case quadratic cost in terms of the size of the biedged graph. Below we consider ways to prune the set of snarls by using their natural nesting relationships to create a hierarchy of snarls that is at most linear in the size of the biedged graph.

### 2.6 Ultrabubbles and Cactus Trees

Given Theorem 1, to determine the ultrabubbles in *B*(*D*) it is sufficient to check for each chain and bridge pair if the separated component is acyclic and contains no tips. As snarls can contain each other, to do this efficiently we decompose the problem into a series of smaller independent problems. We use a modification of the cactus graph called a cactus tree. For a cactus graph *C*(*D*) the *cactus tree T*(*D*) is created by, for each simple cycle *S* in *C*(*D*), making a novel *chain* vertex, adding an edge between each vertex in *S* and *x*, and deleting the edges in *S* (Fig. 3(E)). We call each non-chain vertex (a member of the set *V_C_*_(*D*)_) in *T*(*D*) a *net* vertex. For each chain pair {*x*, *y*} in *B*(*D*) the edge in *T*(*D*) connecting the net vertex projected to by *x* and *y* and the chain vertex representing the simple cycle in *C*(*D*) projected to by the black edges incident with *x* and *y* is the chain pair’s *chain edge*. Each bridge edge in *B*(*D*) projects to the other type of edge in *T*(*D*), which connects two net vertices. Each pair of such edges connected by a path of edges connecting chain and net vertices represents a bridge pair. The edges of a cactus tree *T*(*D*) are therefore decomposable into a set of edges representing the chain pairs in *B*(*D*) and a set of edges representing the bridges in *B*(*D*).

A *parent* snarl contains a distinct *child* snarl if the separated component of the child is contained entirely within the separated component of the parent. From the definition it follows that a snarl that is a bridge pair cannot be contained within another a snarl.

For a snarl {*x*, *y*} in *B*(*D*), let *X* be the path in *T*(*D*) connecting *a_T_*_(*D*)_(*x*) and *a_T_*_(*D*)_(*y*). *X* starts and ends with net vertices and alternates between chain and net vertices. If {*x*, *y*} is a chain pair then *X* consists only of the net vertex which both *x* and *y* project to in *T*(*D*). The *net graph Y* for {*x*, *y*} is a biedged graph as follows (Fig. 3(E)):

- The vertices *V_Y_* are the subset of vertices in *B*(*D*) that project to net vertices in *X*.
- The grey edges in *E_Y_* are the subset of grey edges in *E_B_(_D_)* that connect members of *V_Y_*.
- There is a black edge *e* connecting each pair {*x*′, *y*′} of distinct vertices in *V_Y_* not equal to {*x*, *y*} whose incident black edges in *B*(*D*) both project to the same simple cycle in *V_C_*_(*D*)_ and are connected by a path that starts and ends with black edges.

We can use net graphs to determine if snarls are ultrabubbles. The net graph for a snarl {*x*, *y*} is *bridgeless* if it does not contain a vertex other than *x* or *y* without an incident black edge.

#### Lemma 3.

*A snarl* {*x*,*y*} in *B*(*D*) *is an ultrabubble iff its net graph and the net graph of each snarl contained in* {*x*, *y*} *is acyclic and bridgeless.*

*Proof.* IF: If the separated component of {*x*,*y*} is not acyclic then it contains a directed cycle *S*. Let *e* = {*x*′, *y*′} be a grey edge in *S*. By definition *e* is contained within exactly one net graph *X*. If *x*′ is not incident with a black edge in *X* then its incident black edge in *B*(*D*) is a bridge and *x*′ cannot be a member of a directed cycle in *B*(*D*), therefore let {*x*′,*z*′} be the black edge in *X* incident with *x*′. If *S* does not include *z*′ then there must exist a directed walk in *B*(*D*) from 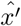,*z*′ that excludes *z*′, but as the black edges incident with *x*′,*z*′ project to the same simple cycle in *C*(*D*), by Lemmas 6 and 11, the deletion of these black edges disconnects *B*(*D*), separating 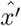 and *y*′, and implying no such directed walk excluding *z*′ be the black edge in *X* incident with *x*′. If *S* does nocan exist. *S* therefore contains *x*′, *y*′, *z*′ and, by the same logic, the node *w*′ in *X* connected by a black edge to *y*′. If *w*′ and *z*′ are not connected by a grey edge then add the nodes in *S* that they are connected to by a grey edge in *X* to this set. Continuing this set extension we must ultimately define a directed cycle in *X*, therefore any cycle *S* in the separated component must define one or more cycles in a net graph. If the separated component contains a tip vertex *x*′ then the incident black edge is by definition a bridge, therefore the net graph containing *x* is not bridgeless.

ONLY IF: Each black edge {*x*′,*y*′} in a net graph *X* represents a portion of a simple cycle in *C*(*D*), there therefore exists a directed path between *x*′ and *y*′ in *B*(*D*) that starts and ends with the black edges incident with *x*′ and *y*′. If there exists a directed cycle *S* in *X* then for each black edge in *S* we can replace it with a corresponding directed path in *B*(*D*) and so define a valid directed cycle in *B*(*D*). If there exists a node *x*′ in *X* not equal to *x* or *y* and without an incident black edge then {*x*′,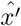 is a bridge edge in *B*(*D*).

It is easily verified that either there exists a tip or a directed cyclic walk in the separable component.

For a chain pair {*x*, *y*} in *B*(*D*) let the *contained chain pairs* be the chain pairs whose chain edges in *T*(*D*) are:

- reachable from *a_T_*_(*D*)_(*x*) without passing through the chain edge of {*x*,*y*} and which,
- on the path from *a_T_*_(*D*)_(*x*), first visit the incident chain vertex and then the incident net vertex.

Similarly for a bridge pair {*x*,*y*} in *B*(*D*) let the *contained chain pairs* be the chain pairs whose chain edges in *T*(*D*) are:

- reachable from a vertex on the path *X* between *a_T_*_(*D*)_(*x*) and *a_T_*_(*D*)_(*y*) without passing through an edge in *X* or the projection in *T*(*D*) of the bridge edges incident with *x* or *y*, and which,
- on the path from a vertex in *X*, first visit the incident chain vertex and then the incident net vertex.

#### Lemma 4.

*For a chain pair or bridge pair* {*x*, *y*} in *B*(*D*) *the set of contained snarls is equal to its contained chain pairs.*

*Proof.* Let *X* be the component of *C*(*D*) containing the projection of *x* and *y* after the deletion of the projection of the black edges incident with *x* and *y*. From Theorem 1 and given that homomorphisms preserve connectedness, it follows that the vertex induced subgraph in *B*(*D*) of vertices that project to a vertex in *X* is the separated component of *x* and *y*. It follows that only snarls whose separated components’ vertex projections are contained in *X* can be contained in {*x*,*y*}. It is easily verified from the definitions that this is equal to the set of the contained chain pairs for {*x*, *y*}.

#### Theorem 2.

*A snarl* {*x*, *y*} *in B*(*D*) *is an ultrabubble iff its net graph and the net graph of each its contained chain pairs is acyclic and bridgeless*.

*Proof.* Follows from Lemmas 3 and 4.

Given Theorem 2 we now sketch an algorithm to compute the set of ultrabubbles for a given bidirected graph *B*(*D*):

1. Calculate *C*(*D*) (e.g. using the algorithm described in [12]).
2. Calculate *T*(*D*).
3. For each chain pair label its chain edge in *T*(*D*) with whether the chain pair’s net graph is acyclic and bridgeless.
4. For each chain pair, traversing from its chain edge in *T*(*D*), use depth first search to determine if its net graph and the net graph of each its contained chain pairs is acyclic and bridgeless, using the labels of the chain edges, and reporting the chain pair as an ultrabubble if so. (By recording if a chain pair is an ultrabubble as it is visited it is easily verified the complete traversal can be calculated by visiting each chain edge only once).
5. Calculate *D*(*D*).
6. For each vertex *x* in *D*(*D*) incident with exactly two edges let {*x*′, *y*′} be the bridge pair whose members project to *x* in *D*(*D*). (There can be at most |*E_B_*_(*D*)_| – 1 such bridge pairs). Calculate if the net graph and the contained chain pairs of {*x*, *y*} are acyclic and bridgeless, reporting the bridge pair as an ultrabubble if true. (As an element in *T*(*D*) can be contained in at most one such bridge pair the cost for this step is *O*(|*V_B_*_(*D*)_| + |*E_B_*_(*D*)_|) for all such bridge pairs).

The computational complexity of steps 1, 2, 4, 5 and 6 is less than or equal to *O*(|*V_B_*_(*D*)_| + |*E_B_*_(*D*)_|) + |*E_B_*_(*D*)_|). For each chain vertex in step 3 the acyclicity of *n* net graphs is calculated, where *n* is the number of simple cycles incident with the vertex in *C*(*D*). In the worst case, this step has a complexity of *O*(|*E_B_*_(*D*)_||*V_B_*_(*D*)_|), which occurs when all vertices in *B*(*D*) are 3-BEC. The cost of step 3 therefore dominates and the worst case complexity of the entire algorithm is *O*(|*E_B_*_(*D*)_||*V_B_*_(*D*)_|). However, if the size of the largest net subgraph is bounded by a constant (which in practice it likely is) the expected running time will be linear in the size of graph.

### 2.7 Rooted Cactus Trees, Ultrabubbles and Genetic Sites

One particularly attractive feature of superbubbles is that they have a nested containment relationship, so that a digraph is partitioned into a set of top level superbubble components and other graph members not contained in a superbubble component, and each top level superbubble component then contains one or more child superbubbles, forming a tree structure. The situation is more complex for snarls and ultrabubbles, in that the separated component of snarls can overlap (Fig. 4), such that each partially contains the other. To create a properly nested hierarchy of snarls it is therefore necessary to exclude some snarls.

Given Lemma 4, to define a hierarchy of snarls that are chain pairs it is sufficient to pick a chain vertex as the root in each component of the cactus forest (e.g. Fig. 4) and only including the chain pairs contained by the root chain, using the definition of chain pair containment defined above. Note that this naturally orients the cyclic chains of chain pairs, breaking them by the chosen chain edge nearest the root.

**Fig. 4.**
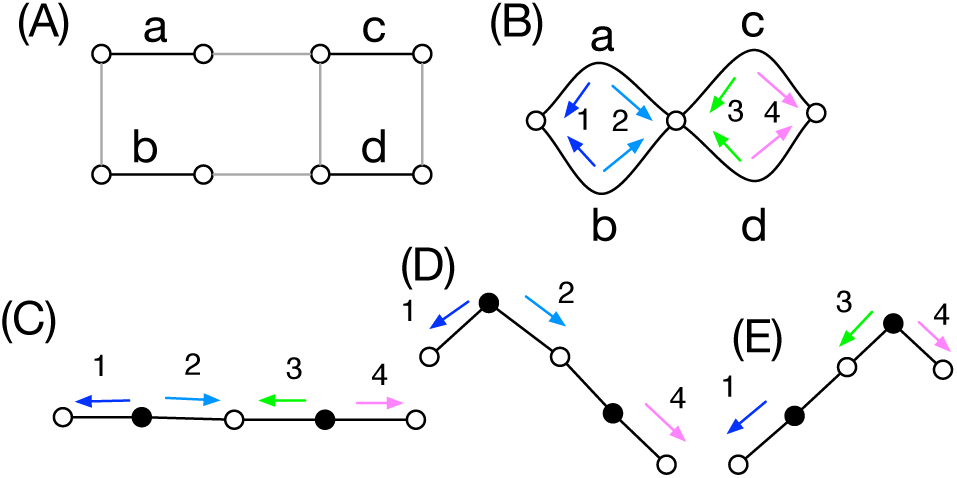
Overlapping snarls. (A) A bidirected graph, its corresponding (B) cactus graph and (C) cactus tree. The snarl numbered 2 contains the snarl numbered 4, similarly the snarl numbered 3 contains the snarl numbered 1. The snarls numbered 2 and 3 overlap. (D-E) Two possible rootings for the cactus tree are shown, each of which defines a properly nested set of snarls.

Snarls that are bridge pairs are not naturally organized hierarchically. However, if the objective is to get a decomposition that contains the maximum number of ultrabubbles then the presence of bridge edges actually simplifies the problem, because a bridge edge cannot be contained within an ultrabubble. Hence if a biedged graph contains bridge edges, we can pick a subset of bridge pairs, and use each bridge pair to define a hierarchy of its contained chain pairs.

One of our motivations for investigating ultrabubbles was to define a decomposition of a bidirected graph representing genome variations into ‘sites’, groupings of paths representing subsequences into ‘alleles’, each representing an alternative at a particular location within a genome. Given a nesting of the ultrabubbles, we can envision that this nesting structure could play a powerful role in decomposing genotyping problems. This is illustrated in Fig. 5, which shows a bridge pair (arrows numbered 1) defining a *top*-*level* ultrabubble. Within this ultrabubble are a series of nested ultrabubbles and chains (*second-level* chain containing snarls numbered 2, 3 and 4 and a *third*-*level* ultrabubble numbered 5). In a genome problem, such a structure can easily arise with nested indels and substitutions. The genotype problem can be viewed as the problem of establishing a consistent genotype of each ultrabubble’s net graph in a coordinated fashion.

**Fig. 5.**
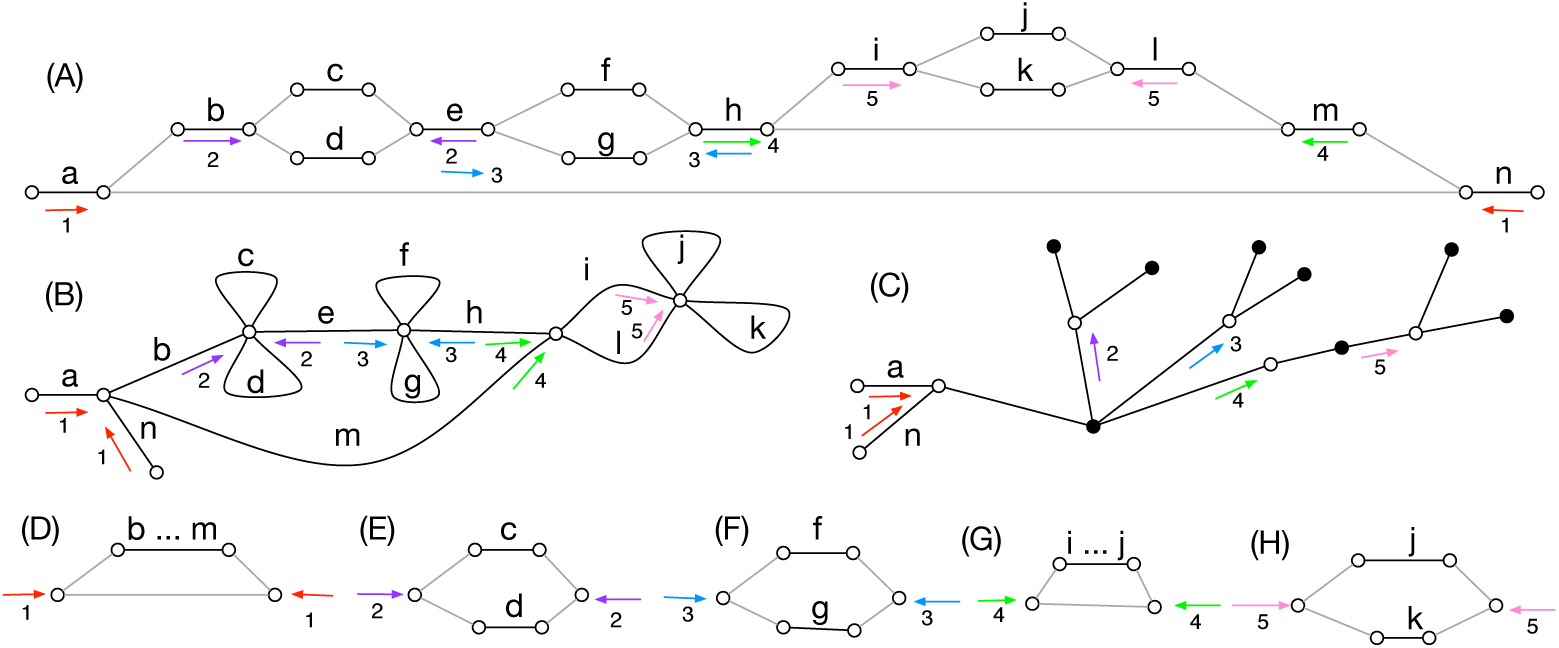
(A) A biedged graph *B*(*D*) with nested ultrabubbles indicated by pairs of numbered arrows. (B) *C*(*D*) for *B*(*D*). (C) *T*(*D*) for *B*(*D*). (D-H) The net graphs for the ultrabubbles in *B*(*D*).

## 3 Results

We implemented the ultrabubbles algorithm described above within the vg software package (http://github.com/vgteam/vg), where it is used to decompose graphs into sites for variant calling. Ultrabubbles can also be computed directly by running `vg stats -u`. To root the decomposition we picked the largest top-level chain, which consists of bridge pairs. Here we present the results of running this decomposition on a graph for human chromosome 1 constructed from the (roughly 6.5 million) variant calls from phase 3 of the 1000 Genomes Project [5]. The graph contained 19, 917, 881 nodes and 26, 782, 661 edges and the runtime was 23 minutes using a maximum of 49G RAM on a single 2.27GHz Intel Xeon core (4 minutes and 30G of RAM were spent loading the graph into memory, a process that can be made an order of magnitude more efficient by switching the implementation to use xg, vg’s succinct representation).

Table 1 shows the relative proportion of each of these structures. The first three rows describe the top-level ultrabubble decomposition, which covers exactly every base in the input graph. The second three rows display the same statistics but for structures that are entirely contained within top-level ultrabubbles or snarls. The remaining rows describe the third and deepest nesting level, which is contained within second level ultrabubbles or snarls. Every base within the graph is part of either a top level chain, ultrabubble or snarl in this decomposition.

**Table 1.** Coverage statistics for the ultrabubble decomposition of the human chromosome 1 variant graph.

| Structure | Nesting Level | Count | Coverage (bp) | Coverage (pct) |
| --- | --- | --- | --- | --- |
| *Chains* | top | 1 | 221,715,143 | 86.60 |
| *Ultrabubbles* | top | 5,554,903 | 12,539,619 | 4.90 |
| *Snarls* | top | 75 | 21,775,387 | 8.50 |
| *Chains* | second | 919 | 20,594,450 | 8.04 |
| *Ultrabubbles* | second | 533,252 | 1,199,777 | 0.47 |
| *Snarls* | second | 0 | 0 | 0 |
| *Chains* | third | 67 | 495 | 0.00 |
| *Ultrabubbles* | third | 694 | 1,623 | 0.00 |
| *Snarls* | third | 0 | 0 | 0 |

Figure 6 shows the size distribution of the top-level ultrabubble and snarl sizes. All but 22 top-level ultrabubbles (totaling 3,251 bases) are 100 bases long or shorter. If we consider such sites “easy” to call, along with top-level chains, then we can assign roughly 91.5% of chromosome 1 into this category. Figure 7 displays three examples of such small ultrabubbles. The remaining 9.5% of cases are found in a small number of relatively large snarls.

**Fig. 6.**
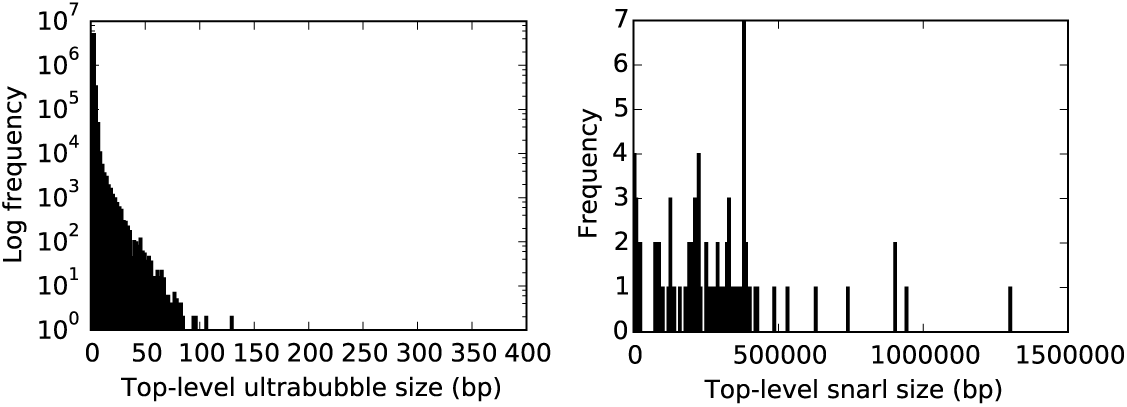
Histograms of top-level ultrabubble and snarl sizes in number of bases, as found in the 1000 Genomes graph for chromosome 1.

**Fig. 7.**
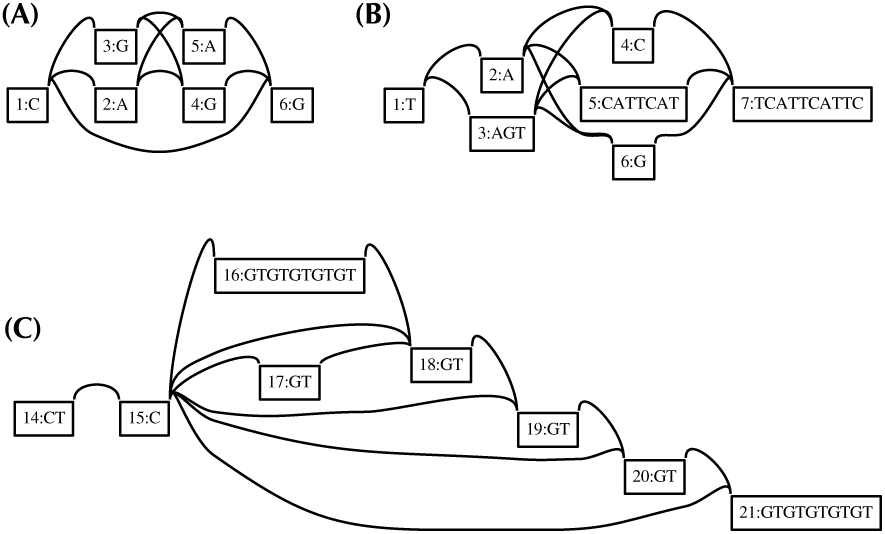
Ultrabubbles found in the 1000 Genomes-derived graph for chromosome 1. (A) Two adjacent SNPs inside a deletion (chr1:209,887,366). (B) A more complex combination of SNP and indel events (chr1:237,977,845). (C) Copy number changes in a GT repeat (chr1:1,200,943).

### 4 Discussion and Conclusion

We have presented a partial decomposition of a bidirected graph into a set of nested snarls and ultrabubbles. We believe this solves an important problem in using graphs for representing arbitrary genetic variations by defining a decomposition that determines sites and alleles.

As the decomposition is only partial, not all elements in a graph will necessarily fit into one of the ultrabubbles. However, we demonstrate that for an existing large library of variation (1000 Genomes) the large majority of sites are either invariant or described by simple, top-level ultrabubbles.

For bases outside of these easy sites it is possible to imagine further subclassification. For example, classifying snarls that contain tips but are acyclic might define a useful class of subgraph common in some subproblems (e.g. sequence assembly). Similarly, characteristic structures representing genomic phenomina, such as inversions and translocations, are imaginable. Beyond our initial investigation, a more thorough evaluation of how much of a graph fits within a snarl, ultrabubble, or one of these more complex structures would be a useful exercise.

As an alternative to further subclassification, in the context of assembly, various error correction algorithms have been proposed to remove graph elements and reduce the complexity of the graph, and therefore correspondingly increase the fraction of the graph that is contained within an ultrabubble structure. We foresee the cactus graph structure providing a useful basis for exploring such algorithms.

## 5 Acknowledgements

This work was supported by the National Human Genome Research Institute of the National Institutes of Health under Award Number 5U54HG007990 and grants from the W.M. Keck foundation and the Simons Foundation. The content is solely the responsibility of the authors and does not necessarily represent the official views of the National Institutes of Health.

## 1 Appendix

### Lemma 5.

*A pair of vertices x*, *y are in the same component of B*(*D*) *iff their projections are in the same component of C*(*D*).

*Proof*. IF: Follows given that by definition no pair of vertices not connected in *B*(*D*) project to the same vertex in *C*(*D*). ONLY IF: Follows given that *a_C_*_(*D*)_ is a graph homomorphism from *B*(*D*) to *C*(*D*) and graph homomorphisms preverse connectedness.

### Lemma 6.

*For a subset of edges X* ⊂ *E_B_*_(*D*)_, *if the removal of the projection of X disconnects C*(*D*), *then the removal of X disconnects B*(*D*).

*Proof*. Follows given that graph homomorphisms preverse connectedness.

### Lemma 7.

*The vertices in C*(*D*) *are the equivalence classes of 3-BEC in B*(*D*).

*Proof*. Each pair of vertices *B*(*D*) that project to the same vertex in *C*(*D*) are either/or-both connected by a path of grey edges (and hence 3-BEC) or connected by at least three black-edge-disjoint paths (using Menger’s theorem).

### Lemma 8.

*A black edge in B*(*D*) *is a bridge edge iff its projection in C*(*D*) *is a bridge edge*.

*Proof*. Let *e* = {*x*,*x*̂} ∈ *E_B_*_(*D*)_

ONLY IF: Suppose *e* is a bridge. As *e* is a bridge the vertices *X*′ reachable from *x* without visiting *x*̂ are black edge connected only by *e* to the vertices *x*′ reachable from *x*′ without visiting *x*. Given Lemma 7, it follows that the projection of *X* and the projection of *X*′ are disjoint, therefore the projection of *e* is a bridge.

IF: Suppose *e* is not a bridge but its projection is. By definition there exists a path in *B*(*D*) from *x* to *x*′ that does not include e. As *a_C_*_(*D*)_(*x*) is a homomorphism, the projection of that path connects *a_C_*_(*D*)_(*x*′) and *a_C_*_(*D*)_(*x*)without traversing *b_C_*_(*D*)_(*x*) implying that it is not a bridge, a contradiction.

### Lemma 9.

*A maximal set of vertices in C*(*D*) *is 2-EC iff the union of its members is a 2-BEC equivalence class of vertices in B*(*D*).

*Proof*. Delete the black bridge edges in *B*(*D*) and the bridge edges in *C*(*D*)′, respectively. Each component is *B*(*D*)′ is, by definition 2-BEC, and similarly each component in *C*(*D*)′ is 2-EC. The proof follows from Lemmas 5 and 8, by showing there exists a bijection between components in *B*(*D*)′ and *C*(*D*)′ such that for each component *X* in *B*(*D*)′ all the vertices in *X* project to vertices in the same component in *C*(*D*)′.

A *cut pair* is a pair of edges whose deletion disconnects the graph.

### Lemma 10.

*A pair of edges in a 2-EC component of a cactus graph is a cut pair iff both edges are contained within the same simple cycle*.

*Proof*. By definition, a 2-EC component of a cactus graph is a set of simple cycles connected by articulation (cut) vertices. It is easily verified that the such a graph is and can only be disconnected by a pair of edges if they occur within one such simple cycle.

### Lemma 11.

*A pair of black edges* (*d*, *e*) *in a 2-BEC component X of B*(*D*) *is a cut pair iff its projection is a cut pair in C*(*D*).

*Proof*. Let *X*′ be a vertex induced subgraph of the projection of *X*. By Lemma 9, *X*′ is a 2-EC component in *C*(*D*).

IF: If the deletion of the projection of *d* and *e* disconnects *X*′ then, using Lemma 6, the deletion of *d* and *e* disconnects *X*.

ONLY IF: If the projection of *d* and *e* are not a cut pair, by the definition of a cactus graph and Lemma 10 the projection of *d* and *e* in *x*′ are each members of two distinct simple cycles. If the projection of *d* (similarly *e*) were a self loop then its endpoints are 3-BEC, implying that after the deletion of *d* and *e* its endpoints remain connected. This is impossible if the deletion of *d* and *e* disconnect the 2-EC component, hence each simple cycle containing the projection of *d* or *e* has length greater than one. For any pair of distinct vertices *x*, *y* in *B*(*D*) that project to the same vertex in *C*(*D*), there exists a path in *B*(*D*) that connects them that excludes their incident black edges, because by Lemma 7 they are 3-BEC, and are therefore either connected by a path of grey edges or, by Menger’s theorem, connected by at least three edge disjoint paths containing black edges. From this observation it is easily verified that the endpoints of *d* (and similarly *e*) must be connected by a path *Y* in *B*(*D*) that includes the black edges that project to the simple cycle containing *d*, in the order of the cycle, and which excludes both *d* and *e*. This implies the endpoints of *d* (similarly *e*) remain connected after the deletion of *d* and *e*, contradicting the claim they are a cut pair.

## 2 Appendix

### Lemma 12.

*Each snarl* {*x*,*y*} in *B*(*D*) *is either a chain pair or bridge pair*.

*Proof*. Using Lemma 5, both *x* and *y* must project to a vertex in the same component of *C*(*D*) as they are connected in *B*(*D*).

Let *d* and *e* be the black edges incident with *x* and *y*, respectively. If *d* is a bridge then *e* must be a bridge, or else by definition *e* connects two vertices in a 2-EC component *X*, the removal of *d* and *e* cannot therefore disconnect *X*, and therefore *y*; and *y*̂, violating the snarl separation criteria. Using Lemma 8, in this case the projections of *d* and *e* must therefore also be bridges. If *d* and *e* are both bridge edges but *x* and *y* do not project to the same vertex in *D*(*D*) (and are therefore not a bridge pair) there exists an intermediate bridge edge *b_D_*_(*D*)_(*z*,*z*̂) on the path between *a_D_*_(*X*)_(*x*) and *a_D_*_(*X*)_(*y*). The deletion *d*, *e* and {*z*, *z*̂} for *B*(*D*) disconnects *B*(*D*) into distinct components, one contains *x* and *z*, one contains *z*̂ and *y*, one contains *x*̂ and one contains *y*̂. This implies {*x*, *z*} and {*z*̂,*y*} each fufill the separation criteria, contradicting the minimality of {*x*,*y*}.

If *d* and *e* are not bridges both must be in the same 2-BEC component or contradict the separation criteria, by the same reasoning as earlier. In this case, Lemma 9 implies both *d* and *e* must project edges in the same 2-EC component in *C*(*D*). Lemmas 10 and 11 further imply they must project to edges in the same simple cycle. If *x* and *y* do not project to the same vertex in *C*(*D*) (and are therefore not a chain pair) then there exists an intermediate black edge *b_C_*_(*D*)_(*z*, *z*̂) on the path between *a_C_*_(*D*)_(*x*) and *a_C_*_(*D*)_(*y*) that excludes *d_D_*_(*D*)_(*x*̂) and *d_D_*_(*D*)_(*y*̂). As with the case that both *d* and *e* were bridge edges, this similarly contradicts the minimality of { *x*, *y*}.

### Lemma 13.

Each chain pair or bridge pair { *x*, *y*} in *B*(*D*) *is a snarl*.

*Proof.* Lemmas 6 and 10 imply that {*x*, *y*} meet the separation criteria. It remains to prove that {*x*, *y*} is minimal. If {*x*, *y*} is not minimal then there must exist an intermediate edge *b_C_*_(*D*)_(*z*, *z*̂) on a path between *a_C_*_(*D*)_(*x*) and *a_C_*_(*D*)_(*y*) that excludes *d_C_*_(*D*)_(*x*̂) and *d_C_*_(*D*)_(*y*̂), and which, using Lemma 12, forms chain or bridge pairs with *a_C_*_(*D*)_(*y*)(*x*) and *a_C_*_(*D*)_(*y*). As *a_C_*_(*D*)_ (*x*) =*a_C_*_(*D*)_(*y*) if {*x*,*y*} is a chain pair, or *a_D_*_(*D*)_ (*x*) = *a_D_*_(*D*)_(*y*) if {*x*,*y*} is a bridge pair, this is clearly impossible.

